# Using deep learning algorithm to surmount drawbacks of statistical method in the procession of identify related genes with cancer

**DOI:** 10.1101/571851

**Authors:** Bohao Zou

## Abstract

Finding key genes which are relative with cancer is the first essential step to understand what has taken place in the tumor cell. At present, the most methods which can discover key genes make a contrast between normal samples and tumor samples and base on the statistical test. However, those methods face on some problems like the insufficient of statistical test in unbalanced samples, defect of only using single data that can not display the holistic situation in tumor cell. For solving those issues, i proposed a innovation method that uses semi-supervised and unsupervised algorithm to discover key genes which are linked with cancer. The genes in the final result list are not only in the double category but with distinct hierarchy and those genes are all detected from diversity data like methylation, gene expression RNA-Seq, exon expression RNA-Seq and so on. At last, for comparing the result of this method and traditional statistical method, i used the conception of information gain ratio to prove the advantage of this deep learning method in mathematical.

## 1. Background & Introduction

As usual, recognition the genes which are different from normal and tumor is essential for knowing the gene function and the role in the developing of tumor. In general, people will firstly use the gene expression RNA-Seq data and statistic method like T-test or Analysis of Variance to calculate the significant of each gene and filter different expression genes (DEGs) between normal samples and tumor samples. For simplified analysis, there are numerous advanced packages in R language like DESeq, edgeR, limma that can assist us to know the different expression genes with cancer. Usually we think, those different expression genes are genes which link with cancer.

However, there are some limitations in those methods for knowing related genes. i) With the complex extent of tumor, the different expression genes may not link with cancer and the non different expression genes may related with tumor. ii) It is a complicate procedure in the occurrence of tumor. only using the single type of tumor data (Gene expression) is not enough to represent the holistic situation in the cell of tumor. iii) T-test, Analysis of Variance or any other statistical methods needs contrast between normal samples and tumor samples. But in the public data sets, the normal samples are significantly less than the tumor samples. It may effect the efficacy of statistic method. iv) If the significant level P-Value of one gene is 0.0101 but the threshold is 0.01. It is hard for us to divide this gene. v) Choosing the threshold of statistical tests has some subjective of human and the quantity of genes will change when select distinctive threshold.

In this work, i proposed a new method by using the newest deep learning algorithm to surmount the drawbacks which i mentioned in the traditional method. I used the Copy number(gene-level), DNA Methylation and Somatic non-silent mutation data to represent the DNA(Genes) mutation level in tumor cells. Used the Gene Expression (RNA-Seq) to represent the RNA expression level in the tumor cells. Finally, used the Exon expression(RNA-Seq) data to represent the expression of protein. For the modeling, i designed a specific convolution neural network (CNN) which i call Diversity Convolution Units Blocks To Simulate Transformer**[1]** With FPN**[2]** NetWork (DCBTF) to classify different tumor samples and then detect genes which are relative with cancer by Class Activation Map**[3]** and K-Means unsupervised learning.

## 2. Data sets

I selected four common cancer types: Breast cancer, Liver cancer, Lung cancer and Stomach cancer. Those data sets are all download from TCGA hub^2^ of UCSC Xean public data hubs^3^. The details information is in the **Table 1**.

**Table 1:**
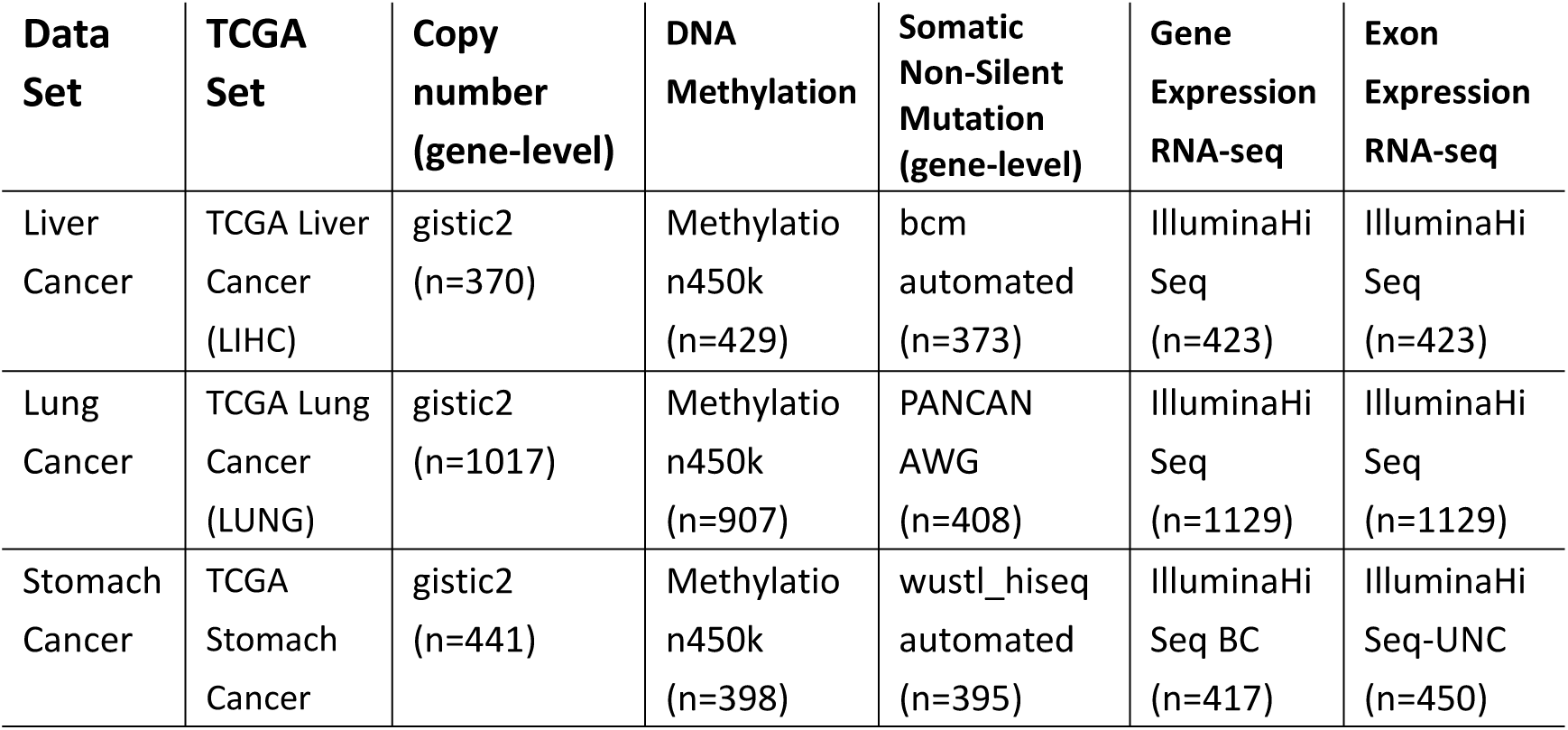
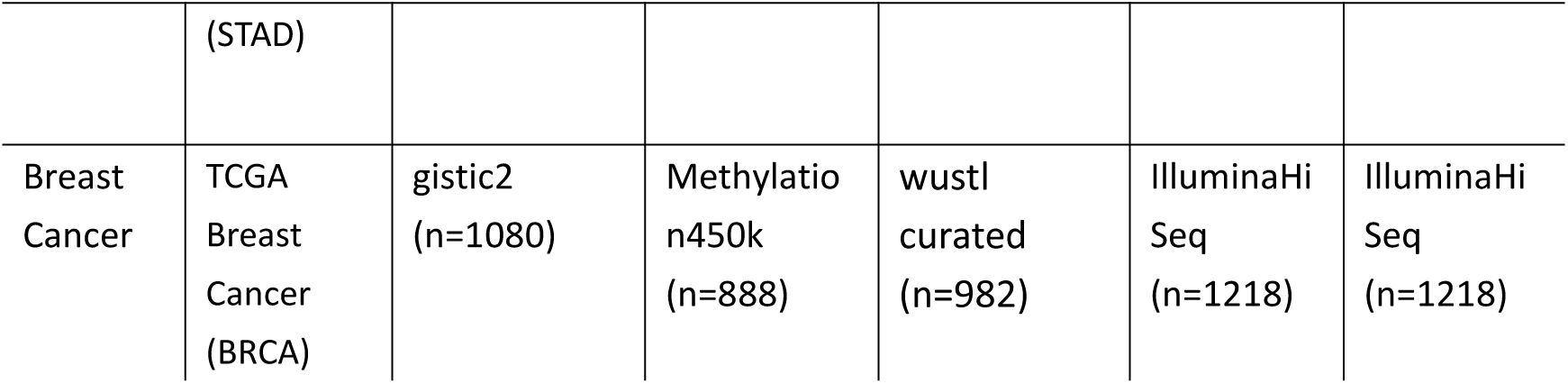
Data sets and its corresponds data.

## 3. Data Processing

In this work, i define the DNA Methylation, Gene Expression RNA-Seq and Exon Expression RNA-seq data as main types data. The Copy Number and Somatic Mutation data as subordinate types. The main types of data are all profile type and the profile type of data can represent one type of information: DNA or RNA or protein independently. The profile type data can represent the integral situation better in the tumor cell with the support of subordinate data. There are six steps in this processing.

Firstly, Due to the data format of the diversity data, we need to split each sample from the data matrix which composes of ids and samples.

Secondly, Deep learning method just learns information from data and dose not need to contrast between normal samples and tumor samples. Another reason is that the quantity of normal, tumor samples are substantial unequally. For solving the problem of unbalance samples. At this step, i filtered the tiny normal samples in each data sets and preserved the tumor samples data.

Thirdly, we need to find samples which contain main data and subordinate data simultaneously. Each sample contains Copy Number, DNA Methylation, Somatic Mutation, Gene Expression RNA-Seq and Exon Expression RNA-Seq, five different types of data. At this step, there are 657 samples in the Breast Cancer, 360 samples in the Liver Cancer, 184 samples in the Lung Cancer and 319 samples in the Stomach Cancer.

Fourthly, we need mapping the id of probe to gene for every sample. The main procession is to deal with the value of each id. If the value with corresponding id is “NA”, i assign 0.0 to this id. if the id has no gene to correspond, i will discard it in my processing. If one gene has many ids to correspond, the value of this gene will be accumulated with all value of id. Finally, there are 24776 gene strings in the data of Copy Number, 26962 gene strings in the data of Methylation450k, 43254 gene strings in the data of Somatic Mutation, 20530 gene strings in the data of Gene Expression RNA-Seq and 22764 gene strings in the data of Exon Expression RNA-Seq.

Fifthly, the input data of the CNN model DCBTF is a multiple channels matrix data like image with “RGB” 3 channels with fixing height and widt. So, in this step, i will trans biology data to this format. Because there are various number of genes in the different types of data, the first thing that we need to do is to uniform the quantity and gene strings between distinctive types data. For the main types data, I seek the intersection gene strings between the main types data and call the set of intersection gene strings as the main genes set. There are 33467 gene string in the main genes set. For the subordinate types data, I will drop the genes in the subordinate types data but not contains in the main genes set. If one gene string appears in the main genes set but the intersection of gene strings of subordinate types data dose not include, i will assign 0.0 to those gene string. At the present, there are 33467 gene string and correspond values in each type of data in one sample. The size of input for training is 192 * 192. Because it is integer times of 2 and facilitates to construct deep learning model. The result of 192 multiply 192 is 36864. The number of 33467 is less than the 36864, so i use 0.0 as place holders for padding those blanks, half of holders add at the head of the image and the other add at the bottom of image. For distinguishing the padding place holders zero and the true zero values, i add a very tiny value to all genes like 1× *e*^−8^. The every pixel in the image represents a gene value in different types data except the padding place holders.

Finally, i normalized all values of genes. I have found that there are extremely huge values in the data and calculated that those values bigger than 10 times of the mean value of one type data in one sample are only 1%. So, i define those values bigger than 10 times of mean value are outlier. I did not discard those abnormal values, but assign its 10 times of mean value. The method of normalizing is Min-Max Normalization.

## 4. Method

As we all know, the performance of classification of CNN has a very strong link with the deep of net, the width of the net, the dense connection between the blocks, the receptive field and the ability of fusing global information.

By the inspiring of Transformer**[1]** encoder network and the state-of-art NLP model BERT**[5]**. I designed the special CNN net named Diversity Convolution Units Blocks To Simulate Transformer With FPN (DCBTF)^4^. The model Transformer-Encoder has a mechanism called Multi-Head Attention that can enhance the dense connection between blocks significantly. So, i selected this model as the basic structure. I have found that the BERT model is only a Multi-Layer Perception model that one single neuron cell has substituted with Transformer-Encoder. So, in this work, i used another more complicate blocks to replace some staple blocks in the Transformer-Encoder model. For increasing the deep and dense connection of the net, i used Dense net**[6]** as one block in my net. For enhancing the width of the net, i also used ResneXT net**[7]** as an part of my net. For the ability of fusing global information, i used Feature Pyramid network**[2]** and average pooling during down sampling to fusing information of extracted features by the backbone net. I used large size filter like 8 × 8 to do convolution and the deep of net for having a bigger receptive field. The batch size in the training may effect the result, for resolving this influence, i used group normalization**[8]**. All method which have mentioned above are all the components of DCBTF net encoder. For the decoder part, i used the transpose convolution to do up sampling operation to substitute the traditional up sampling because i want the the net to learn which location should be focused from the data. Then, i used global average pooling at the bottom of net to transform image which has three dimensions matrix to one dimension vector. Finally, i used only one layer full connection which has no activation function to transform to final output. The nonlinear parts are all in the blocks that contains transpose convolution. The one layer full connection is just to transform the vector to labels quantity which we want net output. This decoder net has substituted the traditional full connection net as the decoder part at the end of normal CNN net. At last, i used L2 regularization to prevent over fitting. The net was constructed by TensorFlow-1.12.0 with GPU version and trained by Nvidia GTX-1080Ti graph card.

For the training step, the initial learning rate is 1× *e*^−5^ and the learning rate would be decay by method of exponential. I used the momentum optimizer to optimize the net. The samples in each cancer type are too small for deep learning. This CNN model should learn the genes expression pattern of one cancer. So, I used 600 breast samples, 324 liver samples, 168 lung samples and 288 stomach samples for training. For the testing set, i used 57 breast samples, 36 liver samples, 16 lung samples and 31 stomach samples for testing. All the training and testing samples are segmented randomly and the proportion of testing samples is around 10% in the total samples. Finally, the accuracy in testing set of breast cancer is : 0.9825, in liver cancer is: 1.0, in lung cancer is: 1.0 and in stomach cancer is: 1.0. The average accuracy is 99.56%. For decreasing the effect of distinction between samples, i would predict training and testing samples through this classification. If one sample predicted error, it would be excluded in next step analysis. There are only 1 sample excluded in all samples. Finally, there are 656 samples in breast cancer, 360 samples in liver cancer, 184 samples in lung cancer and 319 samples in stomach cancer. I define that those samples are filtered samples.

How do we know which location does the net focus on? With the method of **[3]**, we can get an Class Activation Mapping (CAM). This one channel matrix can tell me which location does the net focus on. Some visualization examples are at **Figure 2**.

**Figure 1:**
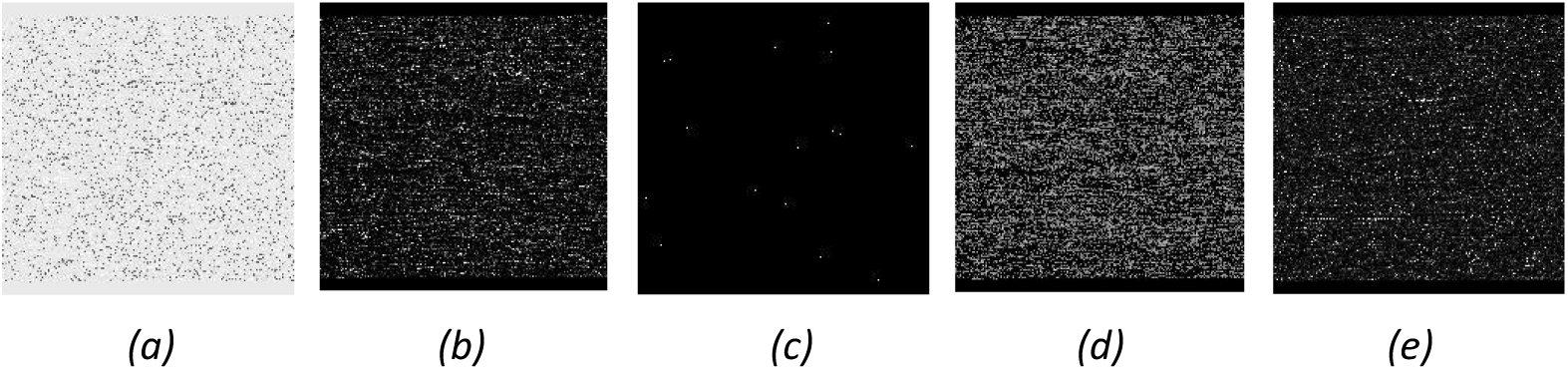
One example of the result of data processing. The sample id in TCGA is TCGA-A1-A0SE-01(Breast cancer sample). **(a)** Copy Number data. **(b)** DNA Methylation data. **(c)** Somatic Mutation data. **(d)** Gene Expression RNA-Seq data. **(e)** Exon Expression RNA-Seq data.

**Figure 2:**
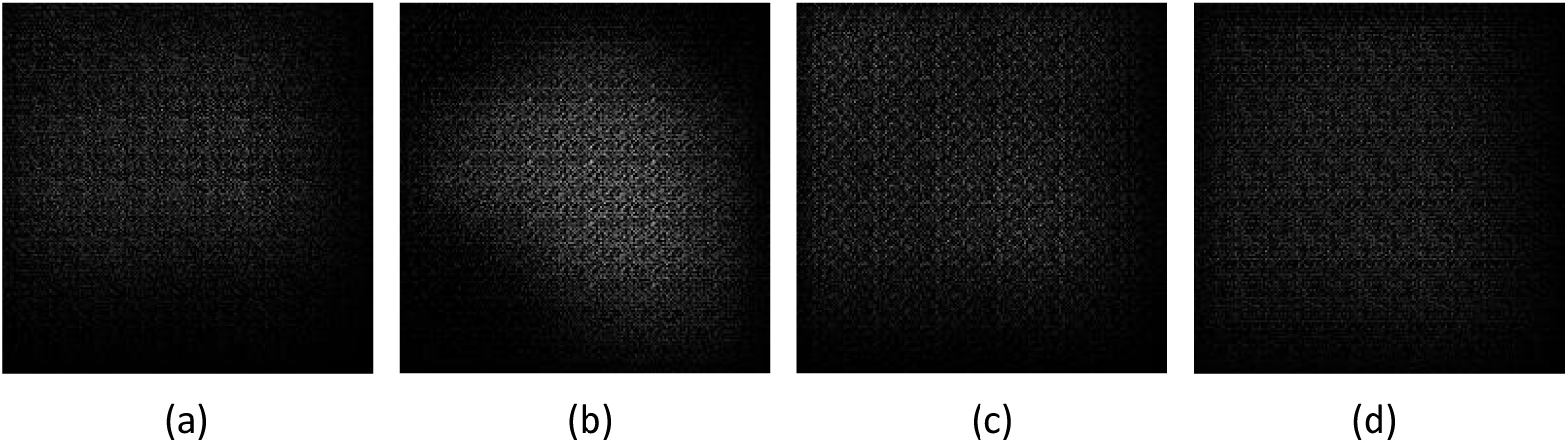
The class activation mapping image of different cancer. (a) Breast Cancer (b) Liver Cancer (c) Lung Cancer (d) Stomach Cancer

How can we separate the related genes? As we all know, the activation location always has higher absolute values and prodigious non-activation location are in lower value interval. The evidence is at **Figure 3**. A numerous values a in the zero neighbourhood.

**Figure 3:**
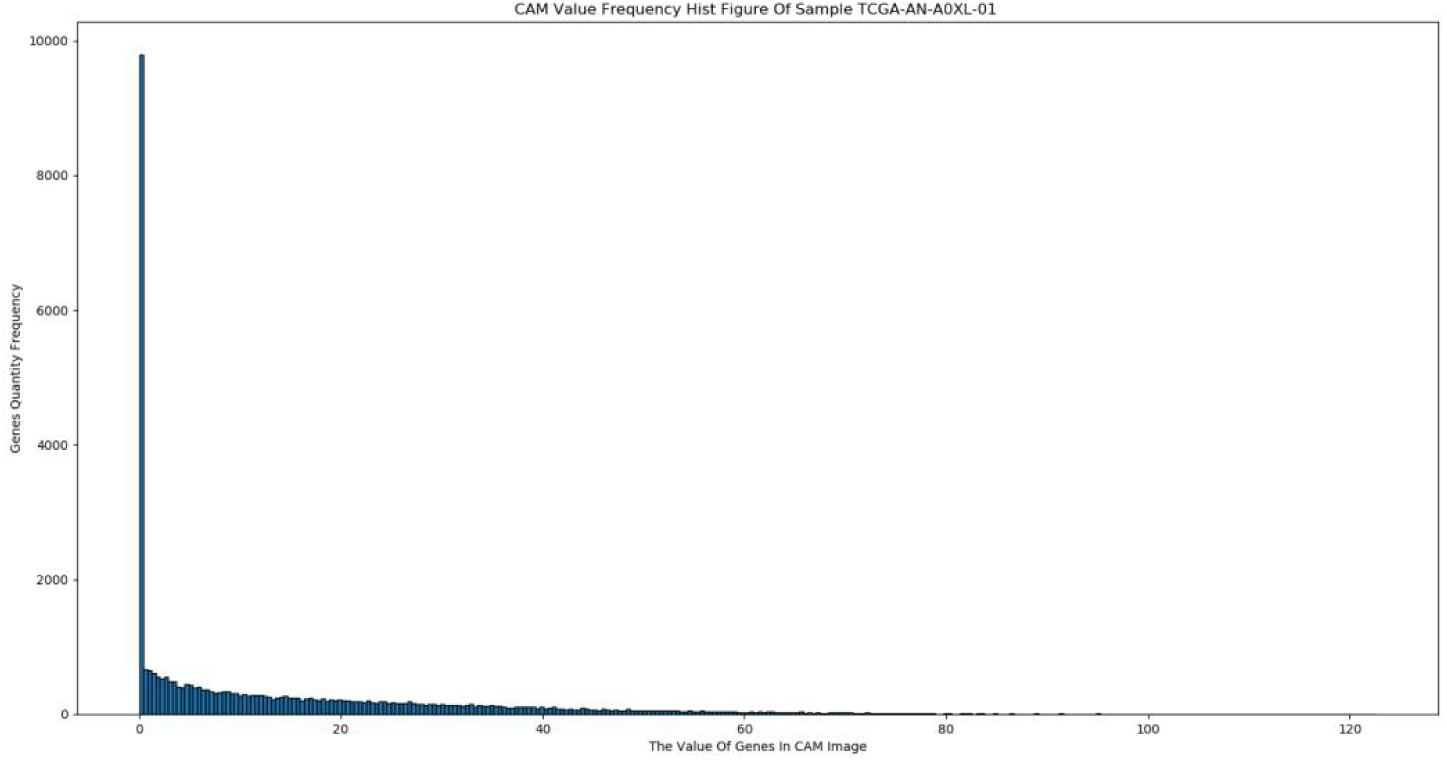
The probability distributions of CAM matrix of sample TCGA-AN-A0XL-01.

Because those high absolute values, the net can discriminate the distinction between different samples. So, at this step, we can pick out those high values with some unsupervised clustering method like K-Means. If we cluster those value into double classes, it must forms a threshold automatically to discriminate the related class or non-related class. If the value of gene is higher than the threshold, it would be clustered into related class. It is same if the value is lower. So, the efficacy of K-Means is poorly enough. Learning from fuzzy theory, i used three classes in the K-Means algorithm, those classes represent non-related genes, probably-related genes and true-related genes respectively. Those classes only link with one sample. The values in CAM matrix were transformed to absolute values for the training in K-Means algorithm. The result of one sample of K-Means is at **Figure 4**. For further analysis, i defined those genes in non-related class gets 0 score, in probably-related class get 1 score, in true-related class get 2 score.

**Figure 4:**
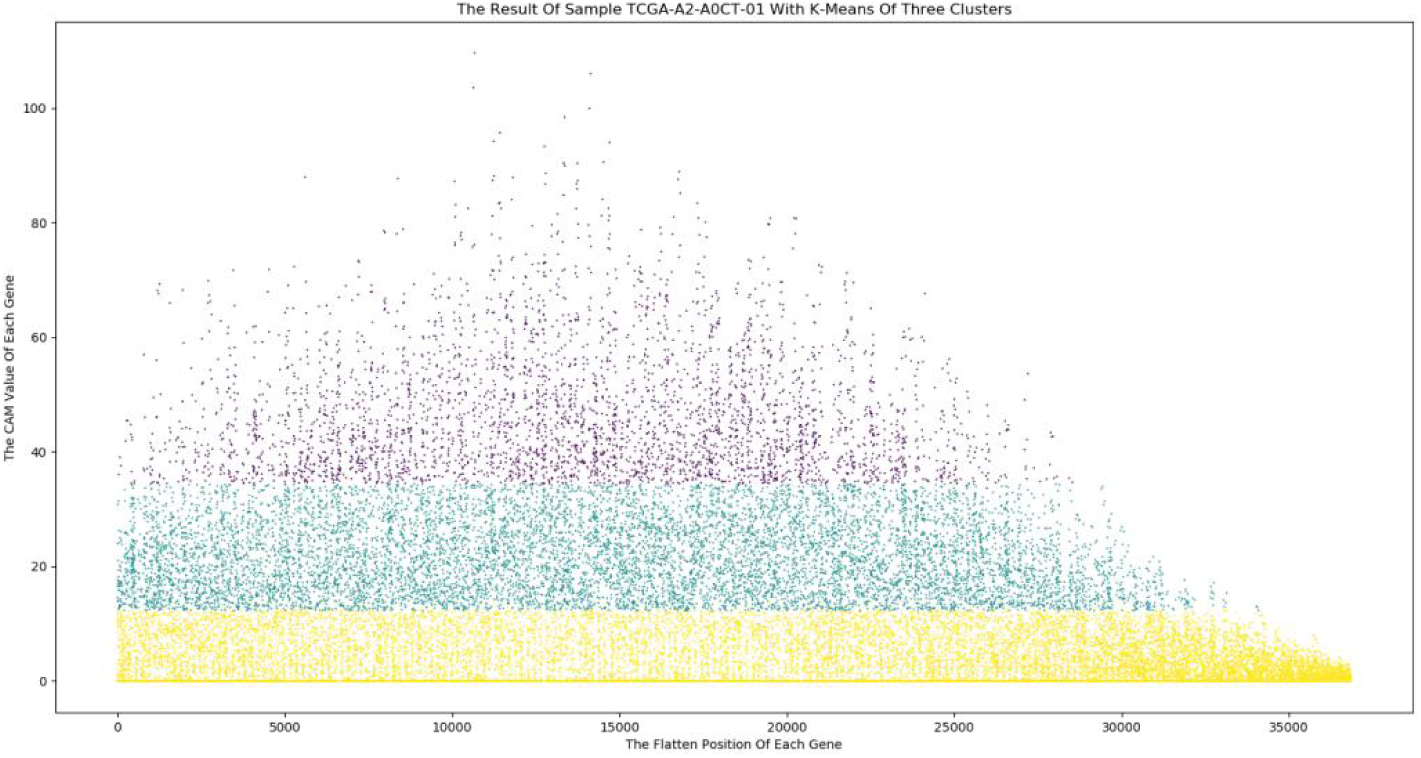
One example result of K-Means. The threshold of non-related and probably-related is approximately at 17 and the threshold of probably-related and true-related is approximately at 38. There are 20314 (60.70%) genes are in the cluster of non-related, 9401(28.09%) genes are in the cluster of probably-related and 3752 (11.21%) genes are in the cluster of true-related.

The values of CAM matrix of each sample have been clustered by K-Means method. Finally, we should statistic the cluster situation of each gene in the whole samples of one type cancer. I would statistic the clustering score and the count number of every gene in the filtered samples at the level of one type of cancer. If one gene in one sample is in the related class (probably or true), i would add 1 to this count number of this gene. All count numbers of every gene are accumulated from zero. At this time, every gene has an clustering score which is added from all samples cluster class score of this gene and the count number value. As the frequency distribution of clustering score of breast cancer as the example, there are 3 peaks in the **Figure 5**. The 3 peaks from left to right represent the number of genes in the score of 0, 656 and 1312. It shows that those non-related and related genes are concentrated in the neighborhood of those peaks. However, there are double huge interval between those three peaks and it is too hard to separate those genes to 3 classes. I will use the fuzzy theory again. Due to there are numerous genes in the interval of peak 0 to peak 656 but less in the interval of peak 656 to peak 1312, it can effect the result if only using 5 classes. I have experiment the situation of having 5 classes. If i used the 5 classes for clustering, there was no center point in the interval of peak 656 to peak 1312. Finally, I would cluster those data to 6 classes by the algorithm of K-Means. The 6 classes represent non-related, probably non-related, weak-related, related, probably strong-related and strong related respectively.

**Figure 5:**
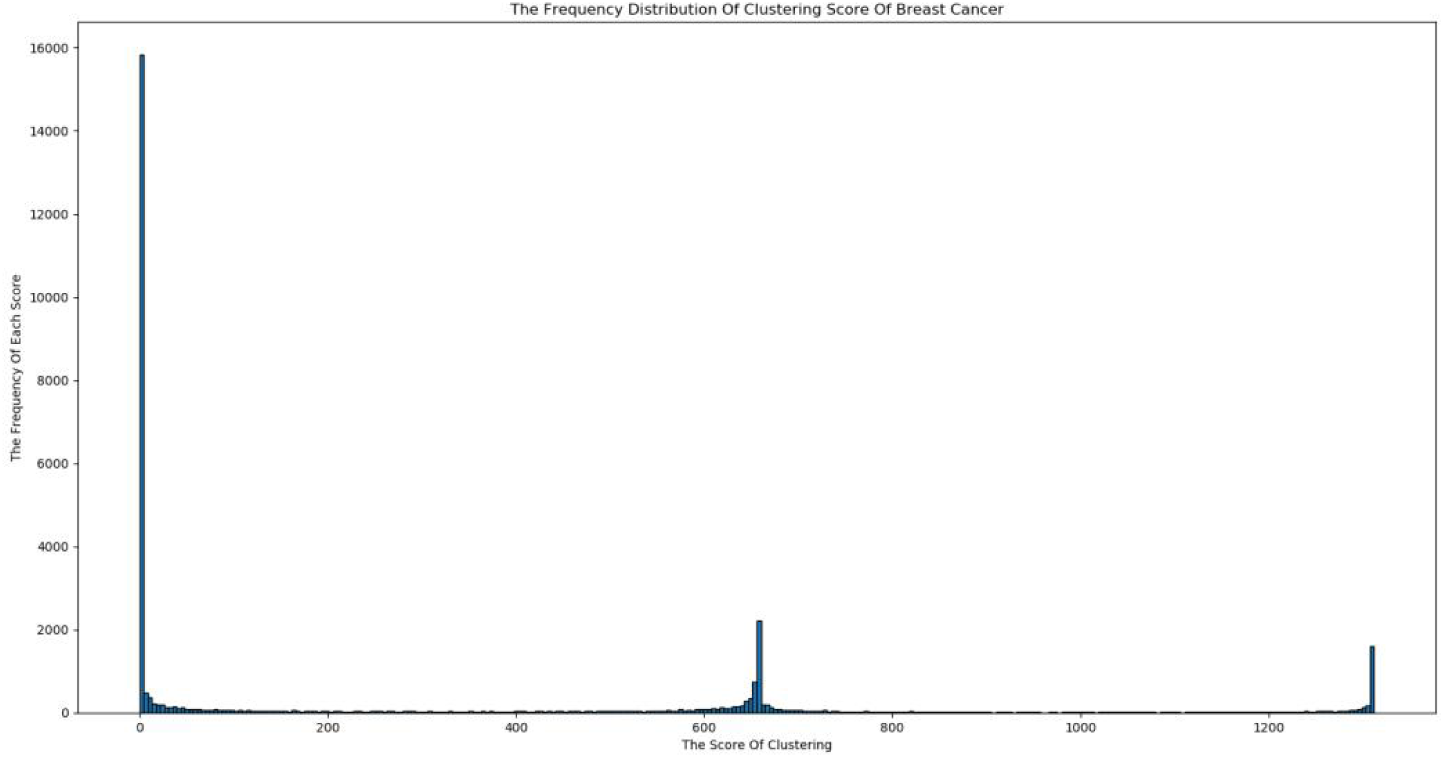
The frequency distribution of clustering score of breast cancer.

With the different classes which clustered by K-Means algorithm, we can assign those genes with different hierarchy. It is may advantages for analysis of down stream.

I also filtered the differential expression genes merely using the data of RNA-Seq with traditional statistical method. The statistical tests which i selected are T-test, Wilcoxon rank-sum test, Levene test and Shapiro-Wilk test. The threshold P-Value of those tests are all 0.01. If the data obey the distribution of normal distribution and pass Levene test, i will use the standard T-test. If the data obey the normal distribution but not pass the Levene test, i will use the corrected T-test. If the data does not obey the normal distribution, i will use the Wilcoxon rank-sum test. Those genes which percolated by statistical test are considered as related genes.

It is hard to compare the difference between those genes which have been discovered by statistical tests with only using RNA-Seq data and this innovation method. So, i have found out an mathematical way to prove the advanced of this innovation method. In my thinking, looking for related genes is like assigning the whole genes into distinct classes. The process of assigning genes is like the split procedure in the algorithm decision tree, assigning data into different sub classes by one selected feature. Moreover, the standard of evaluation uses the information gain or information gain ratio. In this work, we can also use the information gain or information gain ratio to evaluate which split method is better. I have downloaded all genes annotation in the database of Gene Ontology and only used the data in the category of Biological Process. The all genes in this file is data which we want to split and the corresponding IDs of GO are the labels. One gene can correspond multiple IDs. We can calculate the probability of one ID with the frequency of this ID appearing in the all genes annotation file to divide the sum of all IDs frequency number. Furthermore, we can compute the information entropy of the IDs of Biological Process. By the statistical method, we can separate all genes into double classes and also part IDs into double classes simultaneously. By this deep learning method, we assign all genes into six classes and the IDs also be split into six categories. We can calculate the conditional information entropy in those sub sets by using the same method. I will discard the genes if its are not in the all genes annotation file and assign those genes in the annotation file but not in the genes sets that filtered by statistical method or this deep method into non-related class. At this time, we can use information gain or information gain ratio as the norm evaluation. However, the number of classes may influence the result if we use the information gain as the criterion of evaluation. For eliminating this effect, i have used the information gain ratio as the standard finally.

By using those genes in the annotation file as the standard genes, i have leached some genes which are not in the file and generated the final related genes with distinction degrees. Finally, i have done the enrichment analysis in the website Metascape^5^ using those genes in probably strong-related and strong-related category.

## 5. Result

### 5.1 Breast Cancer

The hierarchy result of related genes in the breast cancer is at the **Table 2**.

**Table 2:**
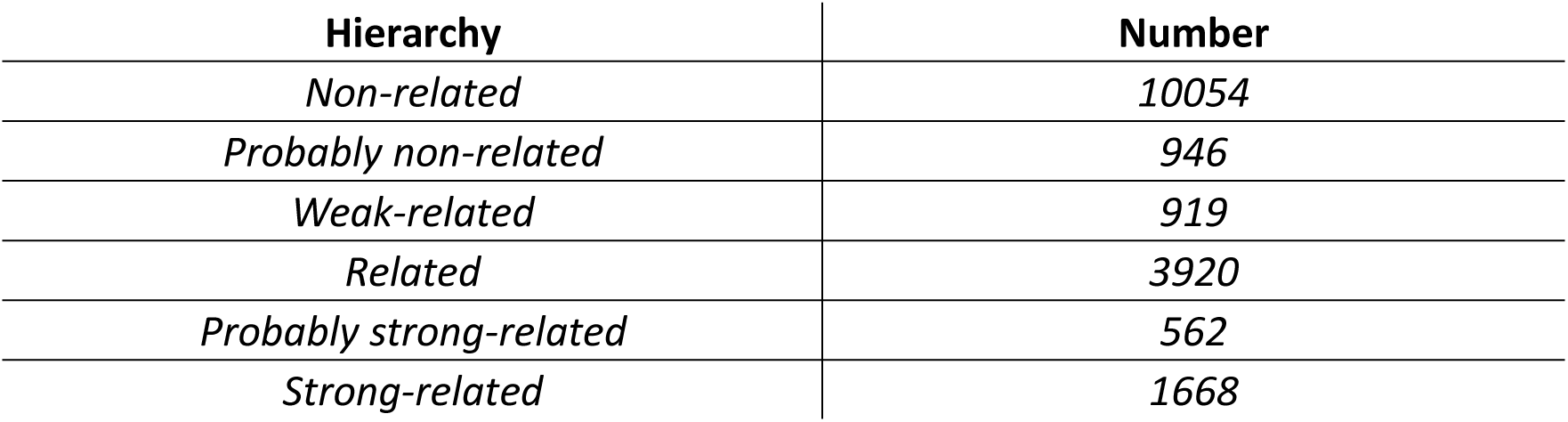
The final result of breast cancer.

| Hierarchy | Number |
| --- | --- |
| *Non-related* | *10054* |
| *Probably non-related* | *946* |
| *Weak-related* | *919* |
| *Related* | *3920* |
| *Probably strong-related* | *562* |
| *Strong-related* | *1668* |

There are 12312 genes which are discovered by statistical tests with the threshold of 0.01 and there are 6873 genes which are all above the level of probably non-related and in the category of related(weak, related, probably strong, strong-related). In this research, the key genes like BRCA1 and BRCA2 are clustered in the categories of probably strong-related and strong-related respectively.

I have drawn a Venn diagram(**Figure 6**) to display the number situation of genes in both different expression class and related category and the genes in different expression class or in related category singly.

**Figure 6:**
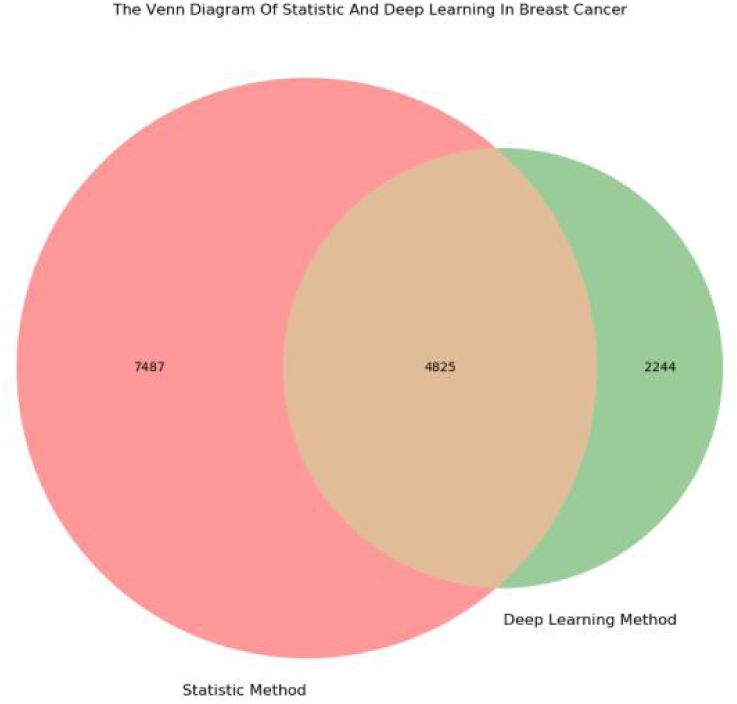
The venn diagram shows the number of genes in each class.

The information entropy of all IDs is 11.7022. The final conditional information entropy that those genes which were divided by statistic method is 11.5954 and the information gain ratio is 0.1273. The final conditional information entropy that those genes which were divided by deep learning method is 11.4382 and the information gain ratio is 0.1461. The improving ratio by using this innovation method is around 14.77%.

I have done enrichment analysis by using those genes in the probably strong-related and strong-related category in breast cancer. The part of enrichment result is at **Figure 7**.

**Figure 7:**
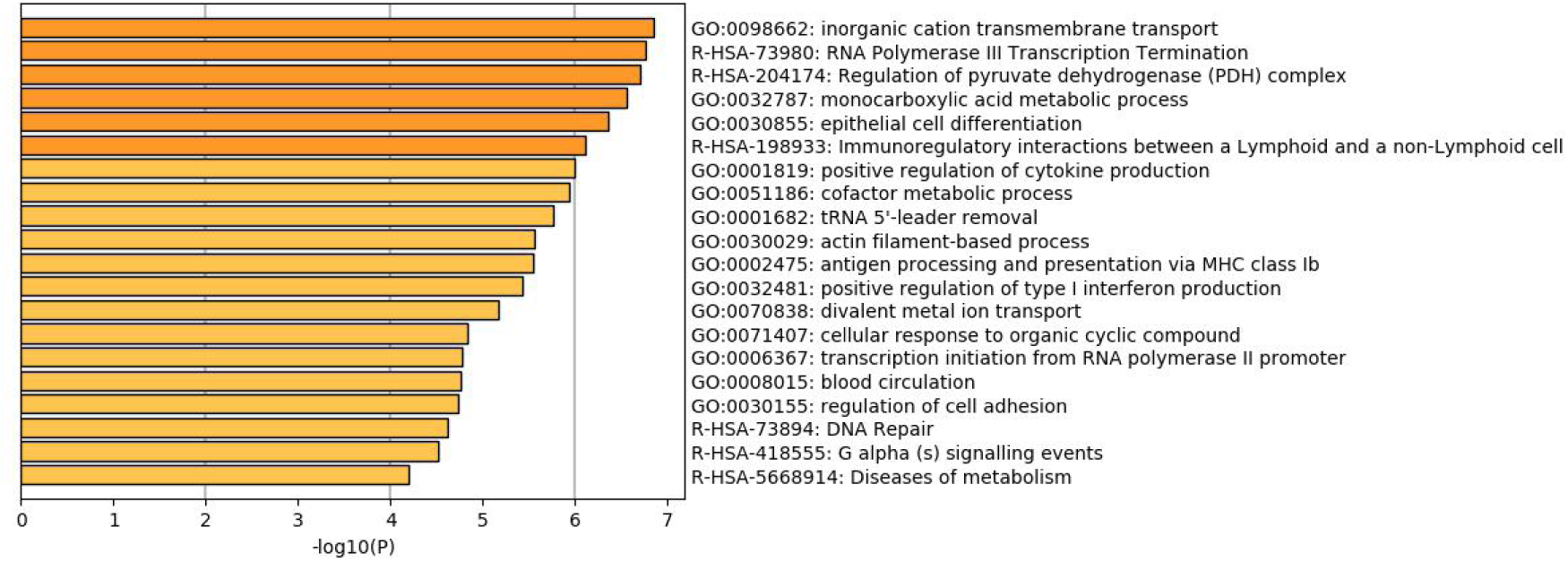
The part result of enrichment analysis in breast cancer.

For further capturing the connection between those terms, the website Metascape selected some enriched terms and display those terms which have the best P-Value as a network plot. If the terms with a similarity > 0.3 are connected by edges. The plot of similarity between those terms with breast cancer is at **Figure 8**.

**Figure 8:**
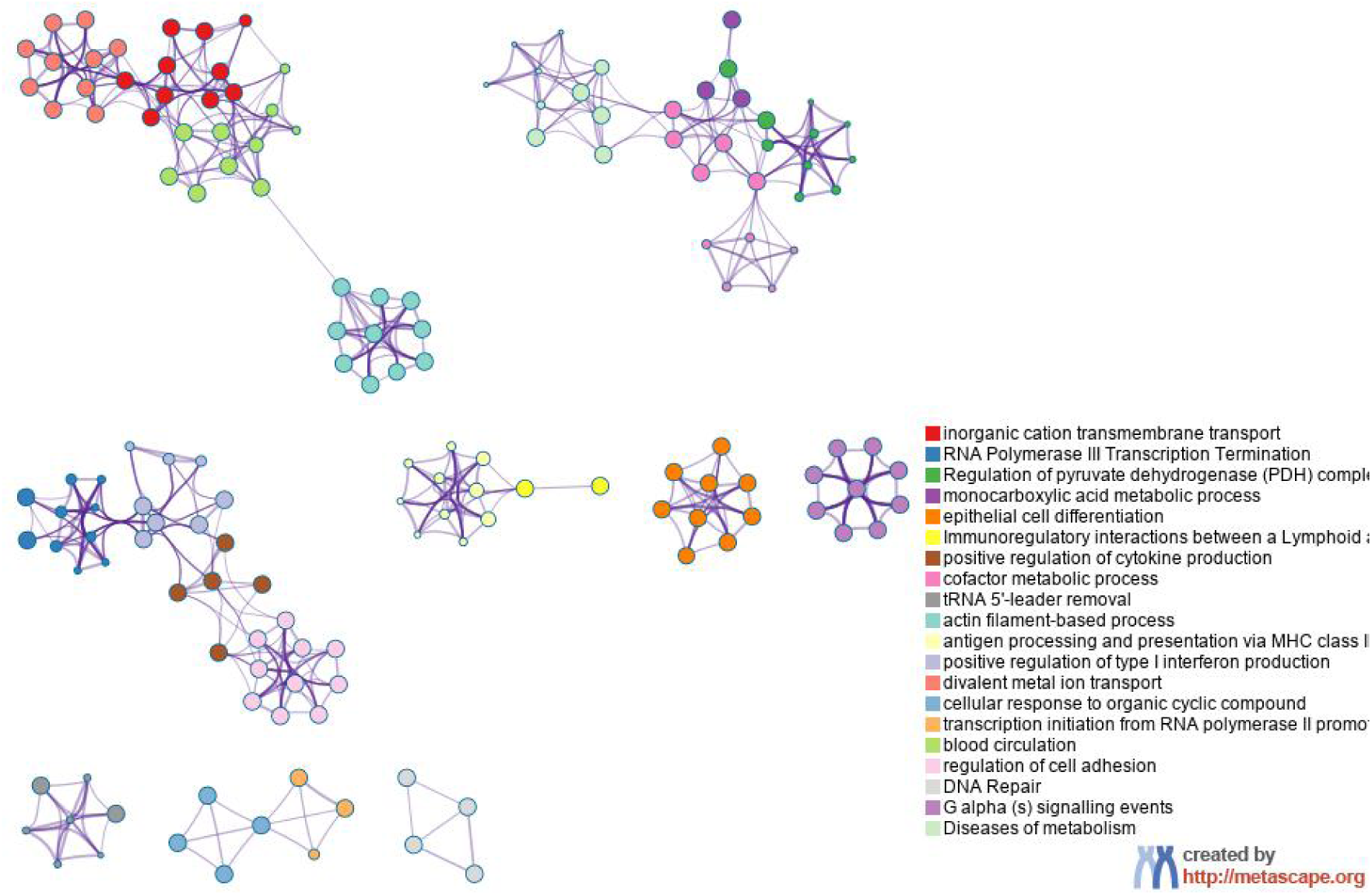
The similarity of terms in breast cancer.

### 5.2 Liver Cancer

The hierarchy result of related genes in the liver cancer is at the **Table 3**.

**Table 3:** The final result of liver cancer.

| Hierarchy | Number |
| --- | --- |
| *Non-related* | *10587* |
| *Probably non-related* | *947* |
| *Weak-related* | *831* |
| *Related* | *3984* |
| *Probably strong-related* | *367* |
| *Strong-related* | *1353* |

There are 10178 genes which are discovered by statistical tests with the threshold of 0.01 and there are 6535 genes which are all above the level of probably non-related and in the category of related(weak, related, probably strong, strong-related). In this research, the key genes like ALDH2 and EPHX1 are clustered in the categories of strong-related and related respectively.

I have drawn a Venn diagram(**Figure 9**) to display the number situation of genes in both different expression class and related category and the genes in different expression class or in related category singly.

**Figure 9:**
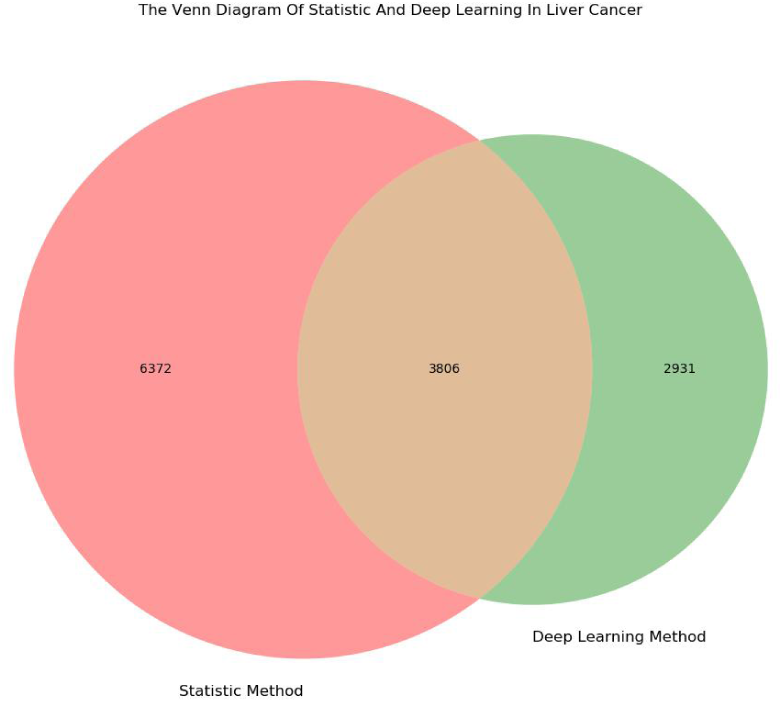
The venn diagram shows the number of genes in each class.

The information entropy of all IDs is 11.7022. The final conditional information entropy that those genes which were divided by statistic method is 11.5974 and the information gain ratio is 0.1082. The final conditional information entropy that those genes which were divided by deep learning method is 11.4659 and the information gain ratio is 0.1450. The improving ratio by using this innovation method is around 34.01%.

I have done enrichment analysis by using those genes in the probably strong-related and strong-related category in liver cancer. The part of enrichment result is at **Figure 10**.

**Figure 10:**
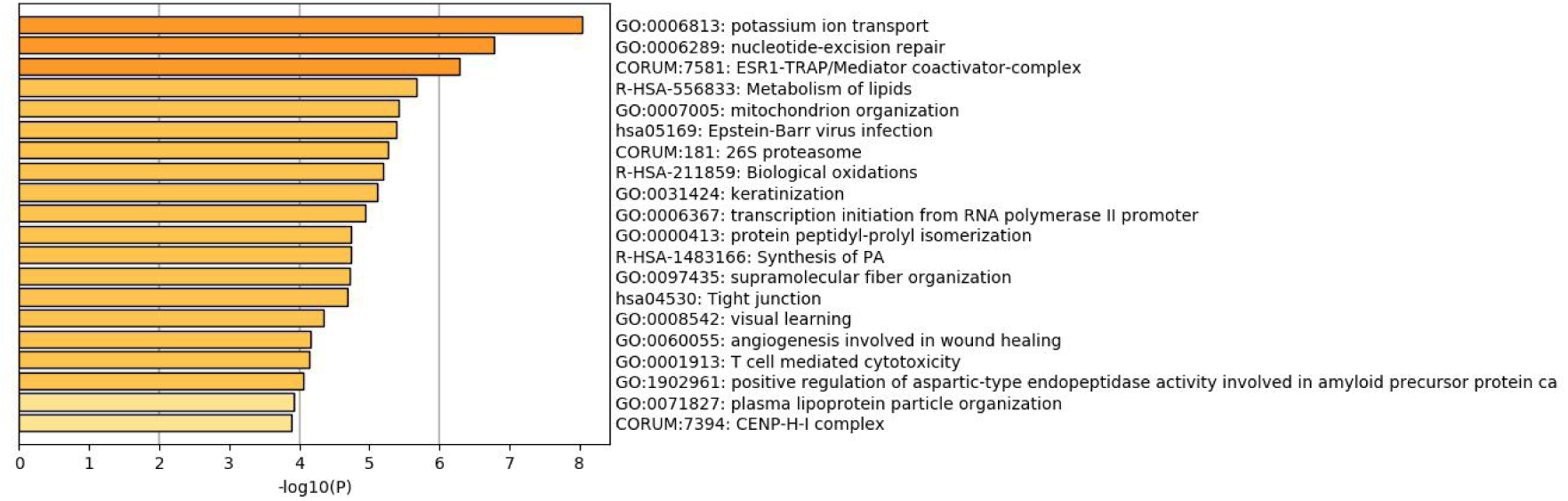
The part result of enrichment analysis in liver cancer.

For further capturing the connection between those terms, the website Metascape selected some enriched terms and display those terms which have the best P-Value as a network plot. If the terms with a similarity > 0.3 are connected by edges. The plot of similarity between those terms with liver cancer is at **Figure 11**.

**Figure 11:**
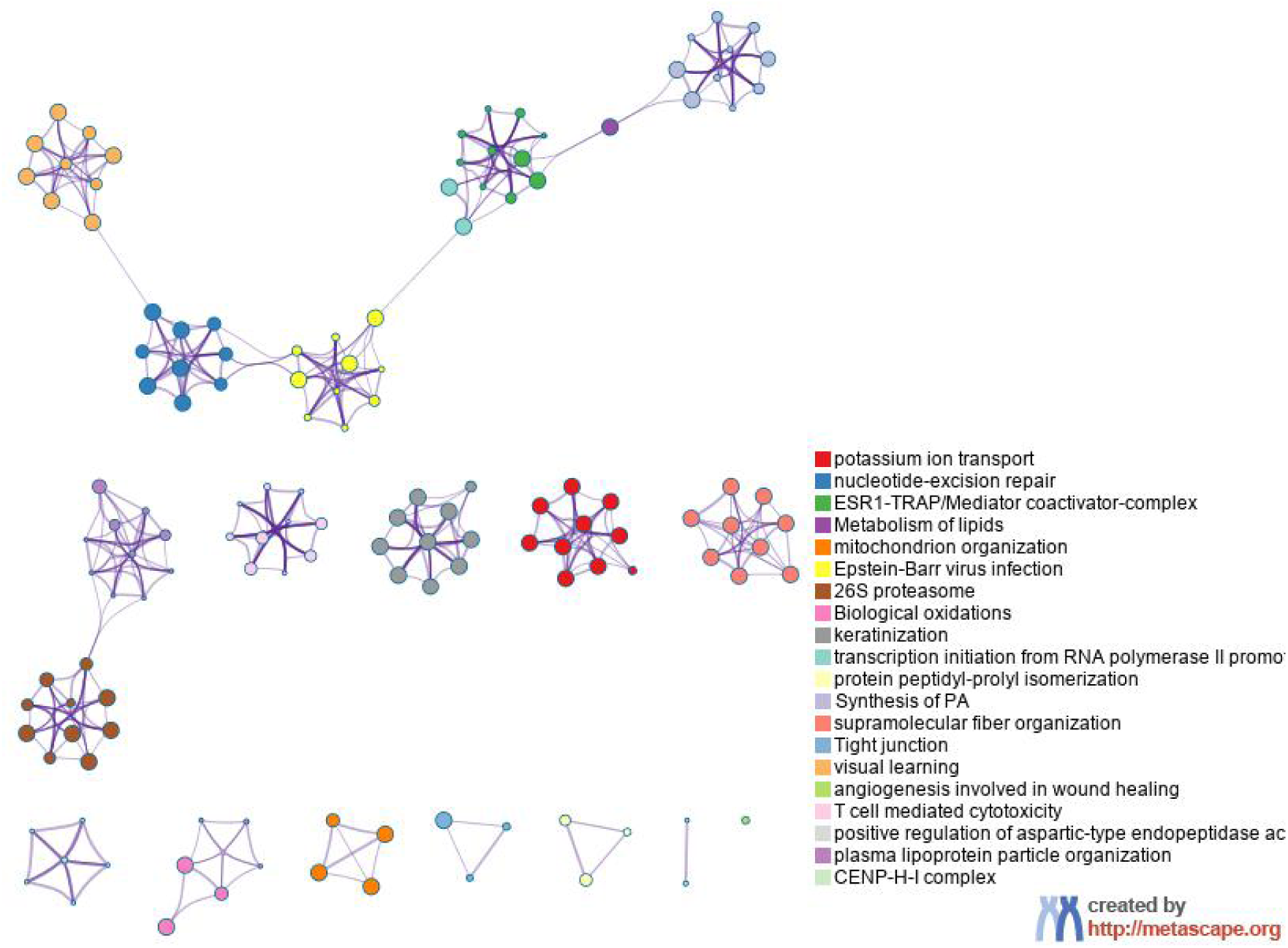
The similarity of terms in liver cancer.

### 5.3 Lung Cancer

The hierarchy result of related genes in the lung cancer is at the **Table 4**.

**Table 4:** The final result of lung cancer.

| Hierarchy | Number |
| --- | --- |
| *Non-related* | 9290 |
| *Probably non-related* | 750 |
| *Weak-related* | 764 |
| *Related* | 4705 |
| *Probably strong-related* | 543 |
| *Strong-related* | 2017 |

There are 12824 genes which are discovered by statistical tests with the threshold of 0.01 and there are 8029 genes which are all above the level of probably non-related and in the category of related(weak, related, probably strong, strong-related). In this research, the key genes like TTF1 and BRAF are clustered in the categories of related and strong-related respectively.

I have drawn a Venn diagram(**Figure 12**) to display the number situation of genes in both different expression class and related category and the genes in different expression class or in related category singly.

**Figure 12:**
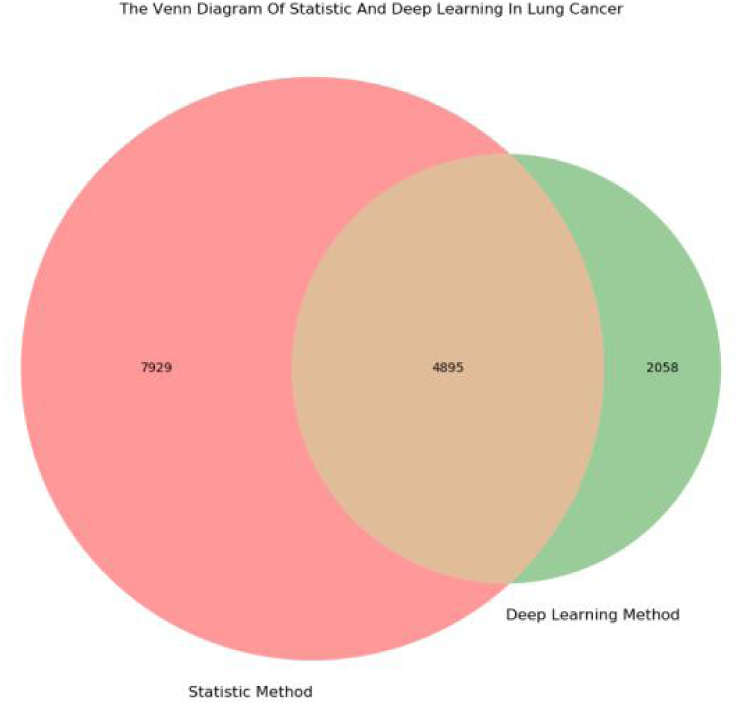
The venn diagram shows the number of genes in each class.

The information entropy of all IDs is 11.7022. The final conditional information entropy that those genes which were divided by statistic method is 11.5968 and the information gain ratio is 0.1308. The final conditional information entropy that those genes which were divided by deep learning method is 11.4674 and the information gain ratio is 0.1437. The improving ratio by using this innovation method is around 9.862%.

I have done enrichment analysis by using those genes in the probably strong-related and strong-related category in lung cancer. The part of enrichment result is at **Figure 13**.

**Figure 13:**
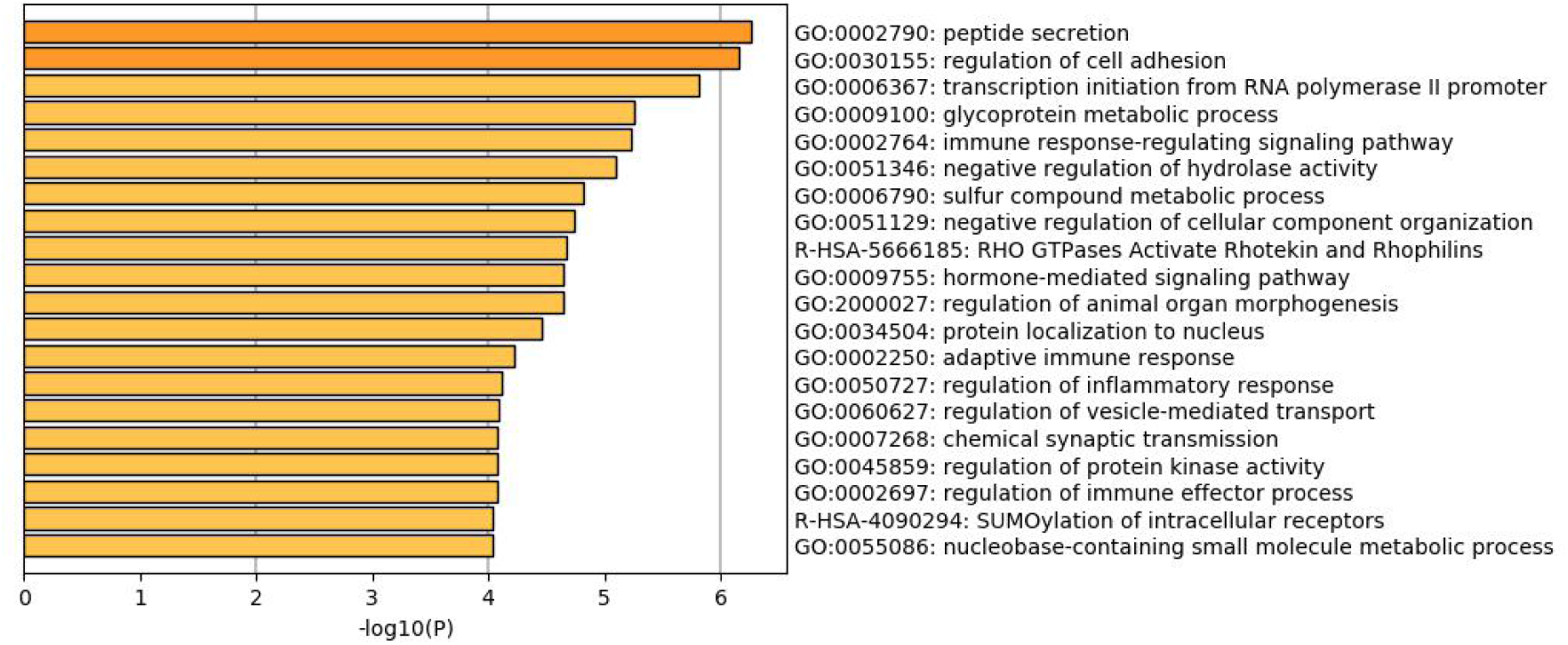
The part result of enrichment analysis in lung cancer.

For further capturing the connection between those terms, the website Metascape selected some enriched terms and display those terms which have the best P-Value as a network plot. If the terms with a similarity > 0.3 are connected by edges. The plot of similarity between those terms with lung cancer is at **Figure 14**.

**Figure 14:**
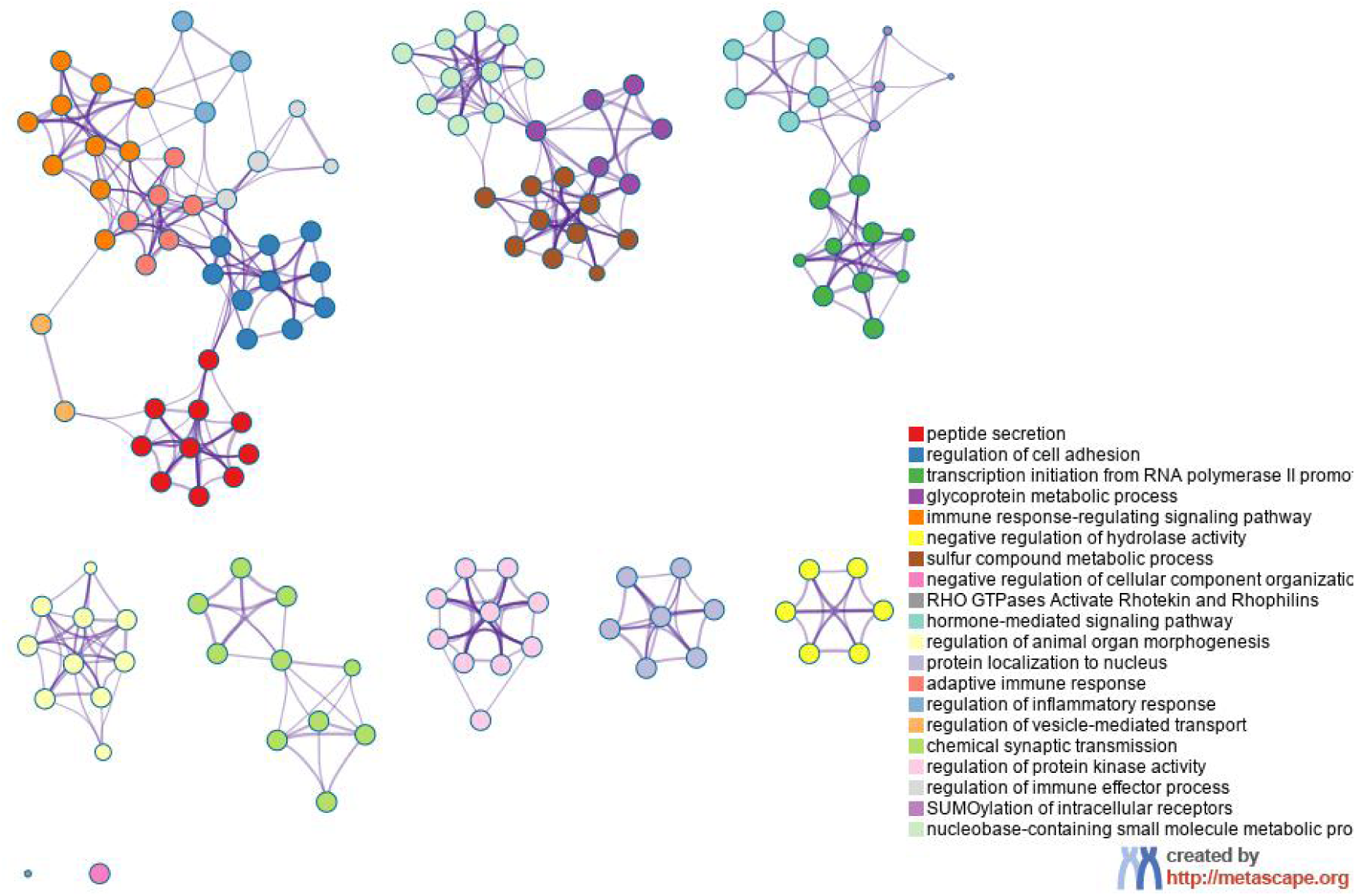
The similarity of terms in lung cancer.

### 5.4 Stomach Cancer

The hierarchy result of related genes in the stomach cancer is at the **Table 5**.

**Table 5:** *The final result of* stomach *cancer*.

| Hierarchy | Number |
| --- | --- |
| *Non-related* | *10271* |
| *Probably non-related* | *414* |
| *Weak-related* | *422* |
| *Related* | *4625* |
| *Probably strong-related* | *227* |
| *Strong-related* | *2110* |

There are 9977 genes which are discovered by statistical tests with the threshold of 0.01 and there are 7384 genes which are all above the level of probably non-related and in the category of related(weak, related, probably strong, strong-related). In this research, the key genes like DCC and IGSF11 are clustered in the categories of related and strong-related respectively.

I have drawn a Venn diagram(**Figure 15**) to display the number situation of genes in both different expression class and related category and the genes in different expression class or in related category singly.

**Figure 15:**
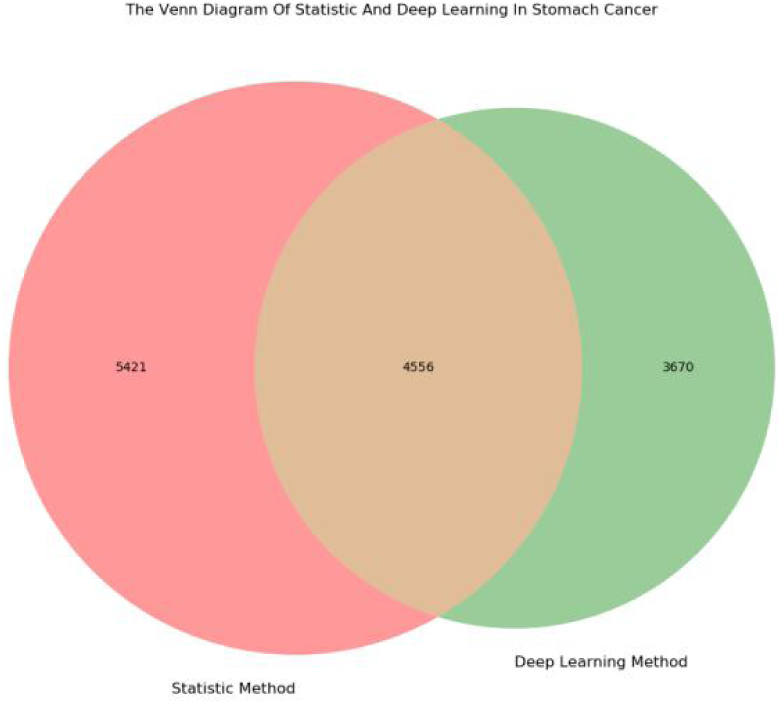
The venn diagram shows the number of genes in each class.

The information entropy of all IDs is 11.7022. The final conditional information entropy that those genes which were divided by statistic method is 11.5883 and the information gain ratio is 0.1155. The final conditional information entropy that those genes which were divided by deep learning method is 11.4808 and the information gain ratio is 0.1381. The improving ratio by using this innovation method is around 19.57%.

I have done enrichment analysis by using those genes in the probably strong-related and strong-related category in stomach cancer. The part of enrichment result is at **Figure 16**.

**Figure 16:**
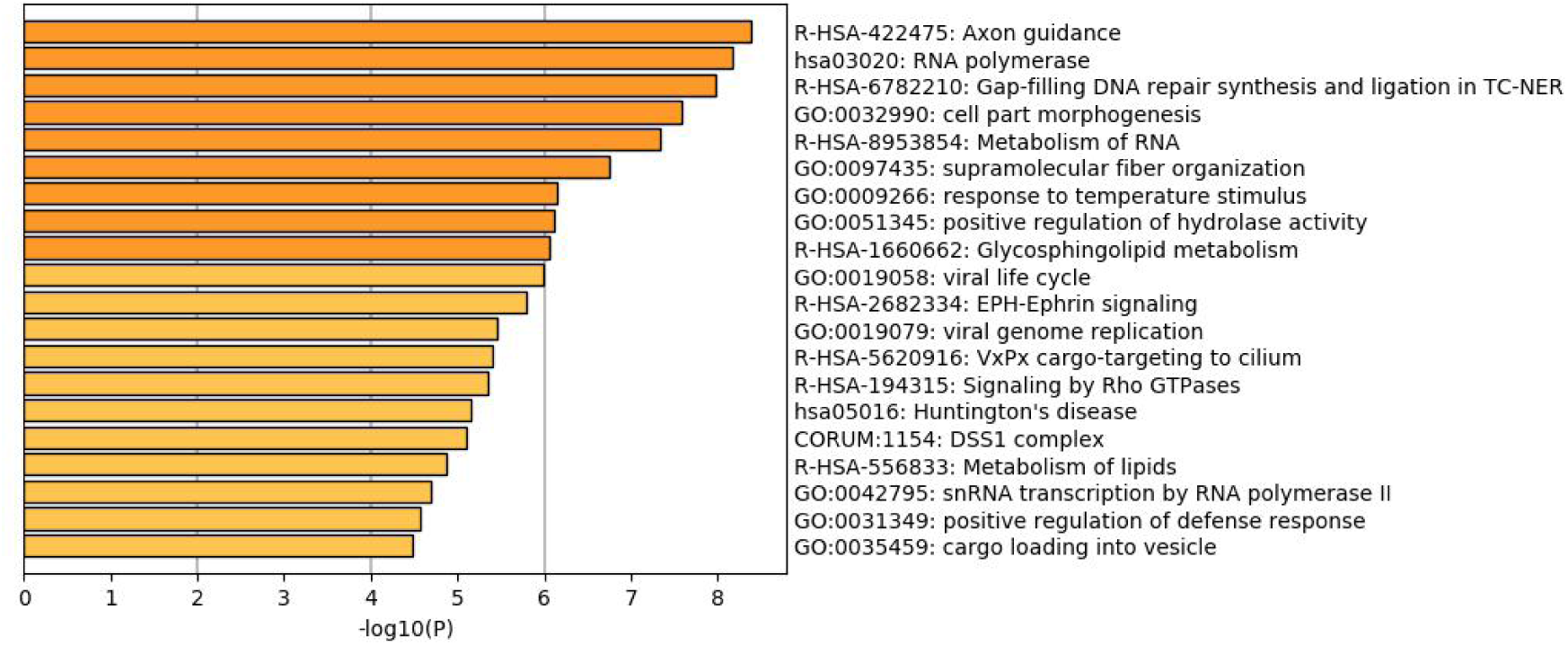
The part result of enrichment analysis in stomach cancer.

For further capturing the connection between those terms, the website Metascape selected some enriched terms and display those terms which have the best P-Value as a network plot. If the terms with a similarity > 0.3 are connected by edges. The plot of similarity between those terms with stomach cancer is at **Figure 17**.

**Figure 17:**
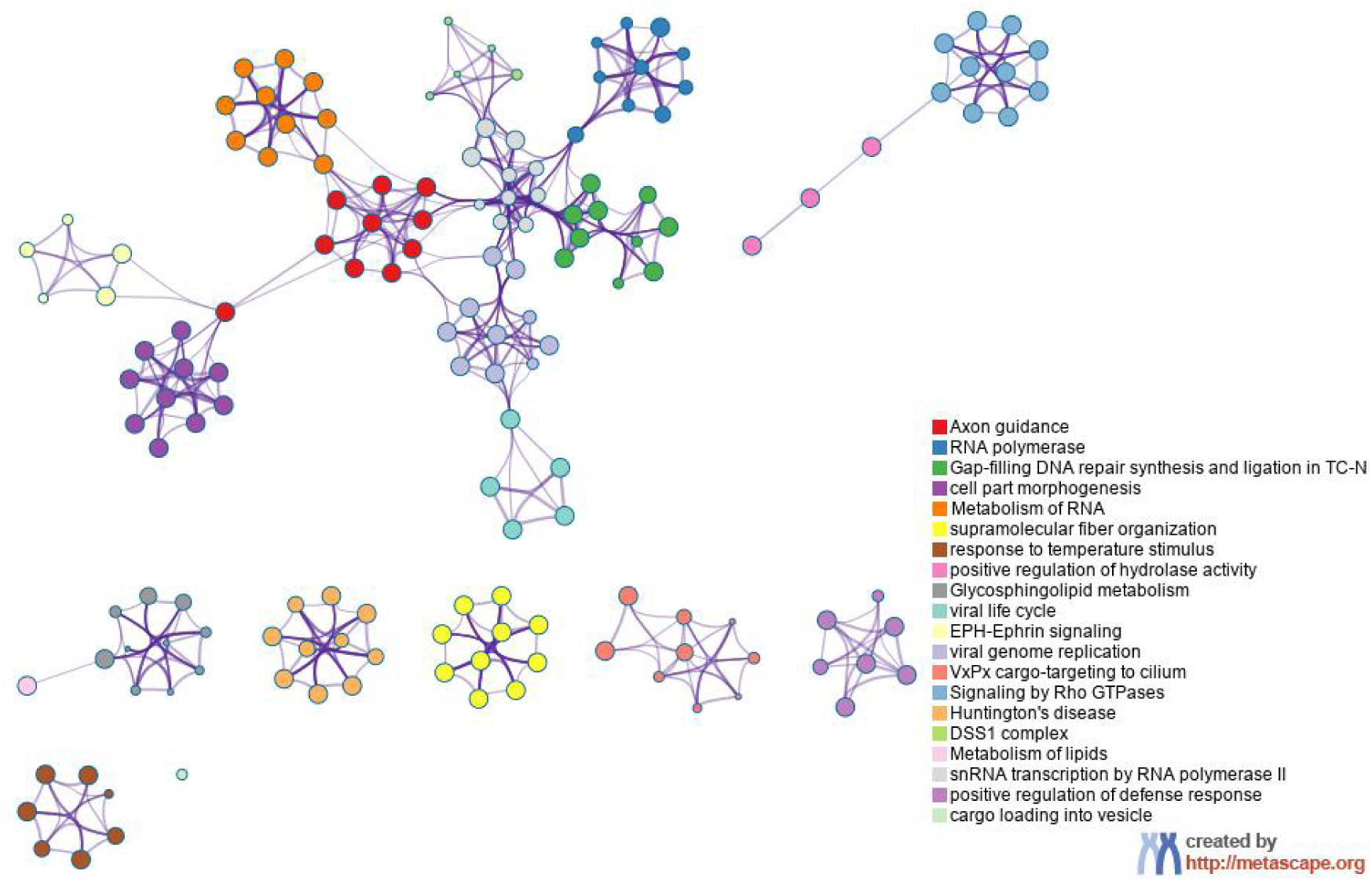
The similarity of terms in stomach cancer.

In the average, the method of deep learning has improved 19.553% than the method of statistical with 0.01 threshold.

With Integrating the information of DNA, RNA and protein in tumor cell, the semi-supervised and unsupervised algorithm application, this method may gives more exact result and has excluded the influence of subjective of human. With the hierarchies of genes, others researchers can select those genes which they are interesting. It is more convenient for down stream analysis.

## 6. Discussion

In this work, i proposed a new method that can surmount some drawbacks in original methods like people may puzzle in choosing threshold, insufficient of statistical test in unbalanced contrasted samples and lack holistic information in tumor cell. This new method discards normal samples to solve the problem of unbalanced samples. Use advanced deep learning semi-supervised algorithm and unsupervised learning to resolve the situation which we all had encountered that confused for selecting threshold. Finally, use multi-dimensional biology data to interpret the circumstance in tumor cell. For the comparing, I have proved that this deep learning method is better than statistical testing in mathematically by using the conception of information gain ratio. However, there is an drawback in this method. With the random selecting initial variables in DCBTF network, The initial variables may effect the final focusing location slightly and this may influence the final clustering result a bit. More straightforward, the result of this deep learning method is not as stable as the result of statistical testing.

## 8. Net Structure

All blocks which will be mentioned next may not depict the channels because in this net, the channels are dynamic and the net is too complicate to describe. If you have interesting to this net, please view my original code in GitHub.

### 8.1 Small-Unit Block

As we all know, in the common model of CNN, we do not only use convolution operation but use the combination of 3 layers, convolution, batch normalization and nonlinear transformation. For reducing the effect of batch size, i use group normalization**[8]** to take place of batch normalization. I choose the PRelu**[9]** as nonlinear transformation, because i want the net to learn negative slope by it self and minimize the influence of gradient disappears and decrease the quantity of death neuron. I define this 3 combine layers as Small-Unit-Convolution-Block(**Figure 18**). This block is the basic block in DBCTF net. It dose not change the size of input, only alter the channels of input to fit some operation like adding.

**Figure 18:**
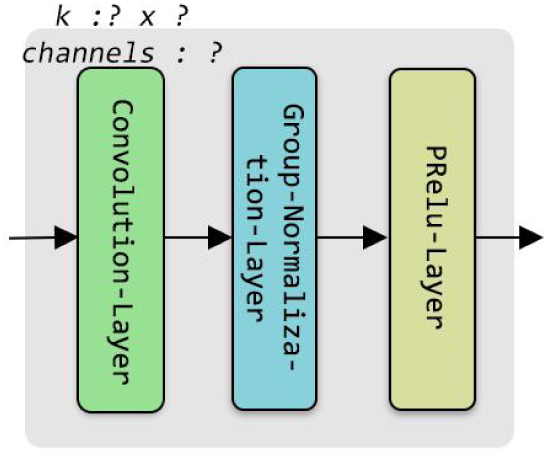
The structure of Small-Unit-Convolution-Block.

### 8.2 Trans-Channels Block

In some situation, the channels of one tensor and another are not same, but we need to add those double tensors or multiply its. At this time, we need to transform those tensors to same channels for those operations. I use the convolution operation to transform channels of one tensor. The convolution operation is an liner operation, it dose not change the traits of original tensor. I define this operation as Trans-Channels-Convolution-Block(**Figure 19**). This is also the basic block in this work. It also dose not change the size of input, only change the channels of input tensor.

**Figure 19:**
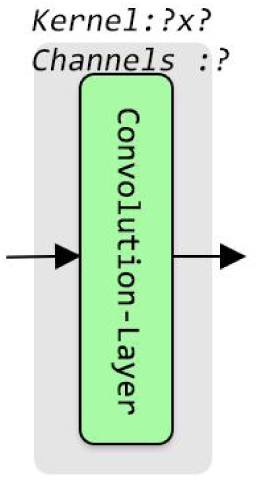
The structure of Trans-Channels-Convolution-Block.

### 8.3 Dense Net Block

Dense net block is used for enhancing the deep of the net. The dense net block is constructed by multiple one dense blocks. In this net, the one dense block establishes with 3 small-unit blocks which have 1×1, 3×3, 1×1 size of kernels respectively. Because the channels of output tensor of one dense block must be same for adding. So, the channels of small-unit blocks in one dense block are out-channels, out-channels divisible 2 and out-channels respectively. For building the dense net, i used 5 one dense blocks in dense net backbone (**Figure 20**).

**Figure 20:**
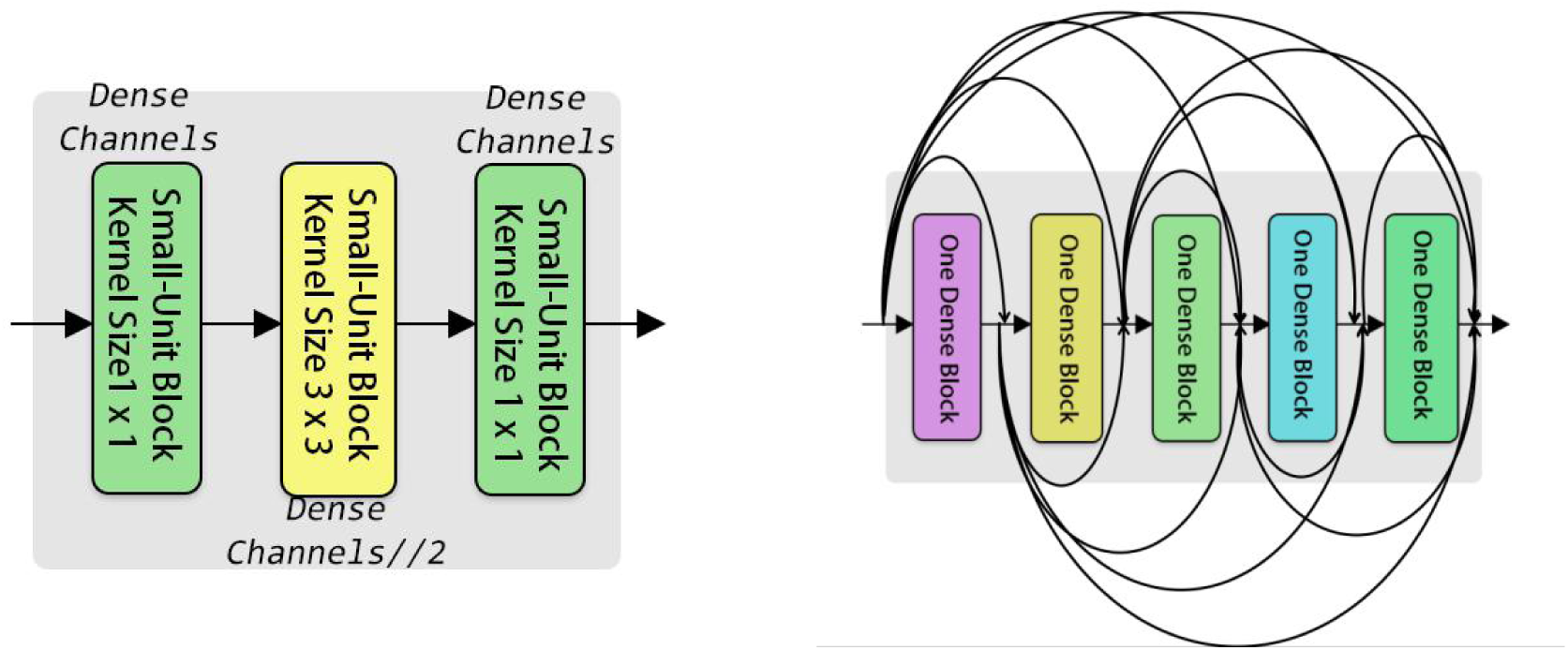
The structure of one dense block (Left) and the structure of dense net backbone (Right). The dense channels in the figure represents the out-channels in passage.

In the dense net block, i also used the mechanism of attention in the image**[10]**. Used global average pooling to down sampling image from [H,W,C] three dimension to [C] one dimension. Then used double full connection layers with C // 16 and C neurons with PRelu activation function to transform channels factors. After then, use up sampling to restore the [C] vector to [H,W,C] three dimensions. Finally, i use sigmoid function to map factors to the interval of zero to one.

The output channels of this dense net block are dynamic. At the end, the details of dense net block is at **Figure 21**.

**Figure 21:**
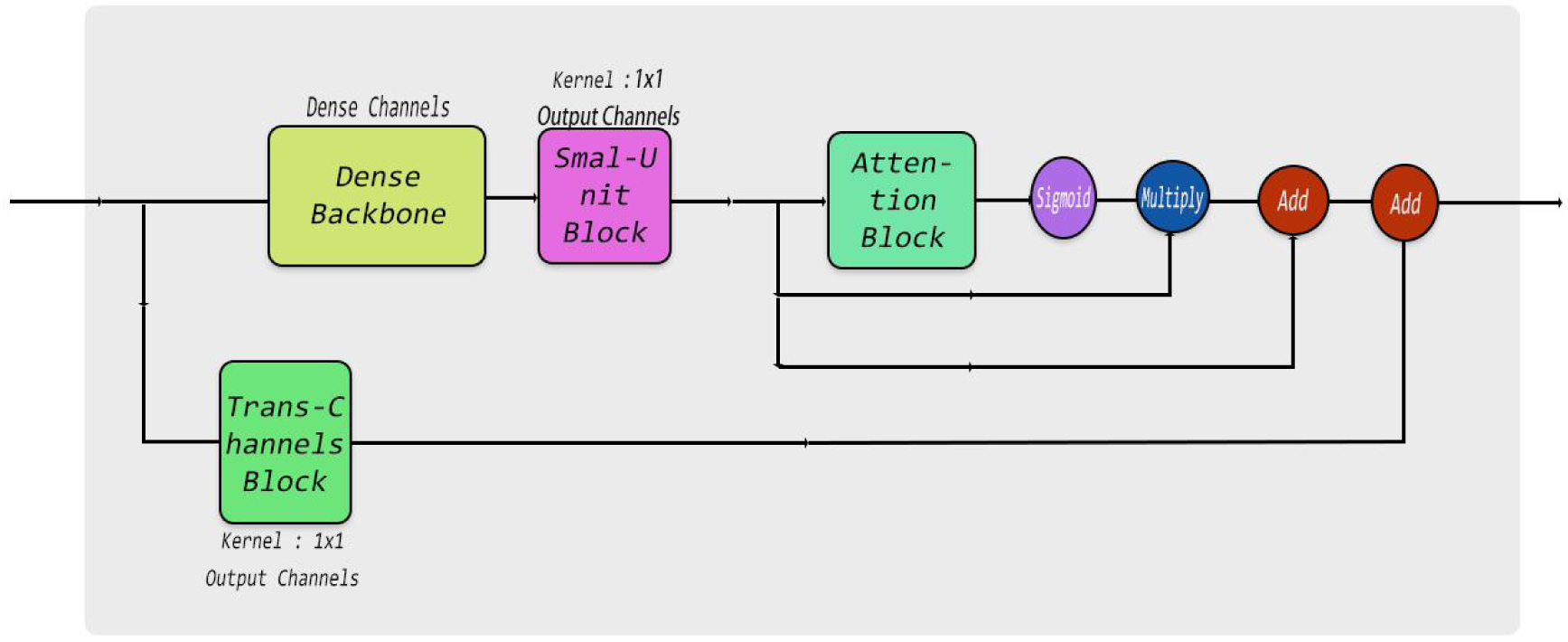
The final structure of dense net block.

### 8.4 ResneXT Net Block

ResneXT net block is used for improving the width of net. In the backbone of ResneXT, it has prodigious parallel blocks with tiny channels. In one parallel blocks, i used three Small-Unit blocks with 1×1, 3×3, 1×1 and 8, 8, out-channels as three Small-Unit blocks kernels and channels respectively. At the end, all the results of parallel blocks would be added to merely one final result. The quantity of parallel blocks are also not stationary. The attention mechanism is the same as dense net block.

The structure of ResneXT net block is similar with dense net block. The structure of ResneXT net is at **Figure 22**.

**Figure 22:**
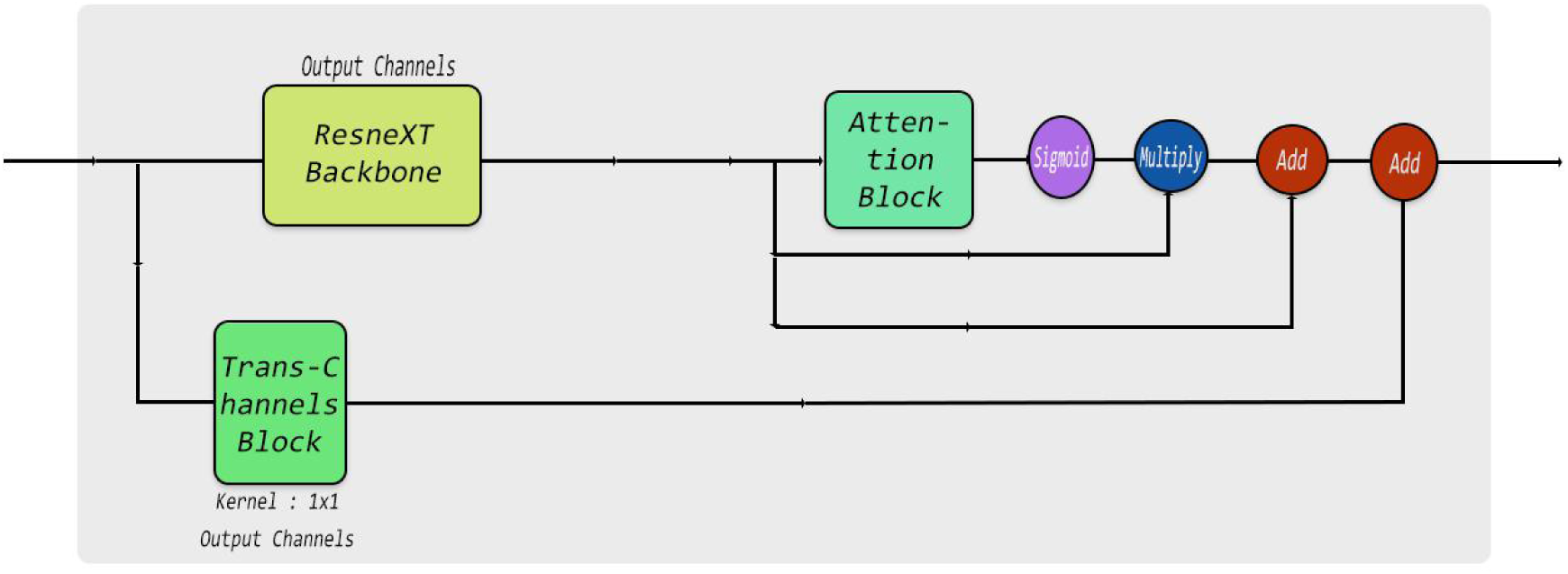
The structure of ResneXT-Net block.

### 8.5 Combine ResneXT net and Dense net block

How to use those advantages of different net structure? I designed the combine block to use those advantages of each net structure. The details of the combine block are at **Figure 23**. There are double scalar variables and have used sigmoid function to convert value to the interval of (0, 1). Those variables can be learned from the data. After that, those variables would be scalar multiplied with the result of ResneXT net block and dense net block respectively. The function of those variables is like an information gate to control how many information can pass the gate. Finally, the information of passing the gate will be added as the terminal output.

**Figure 23:**
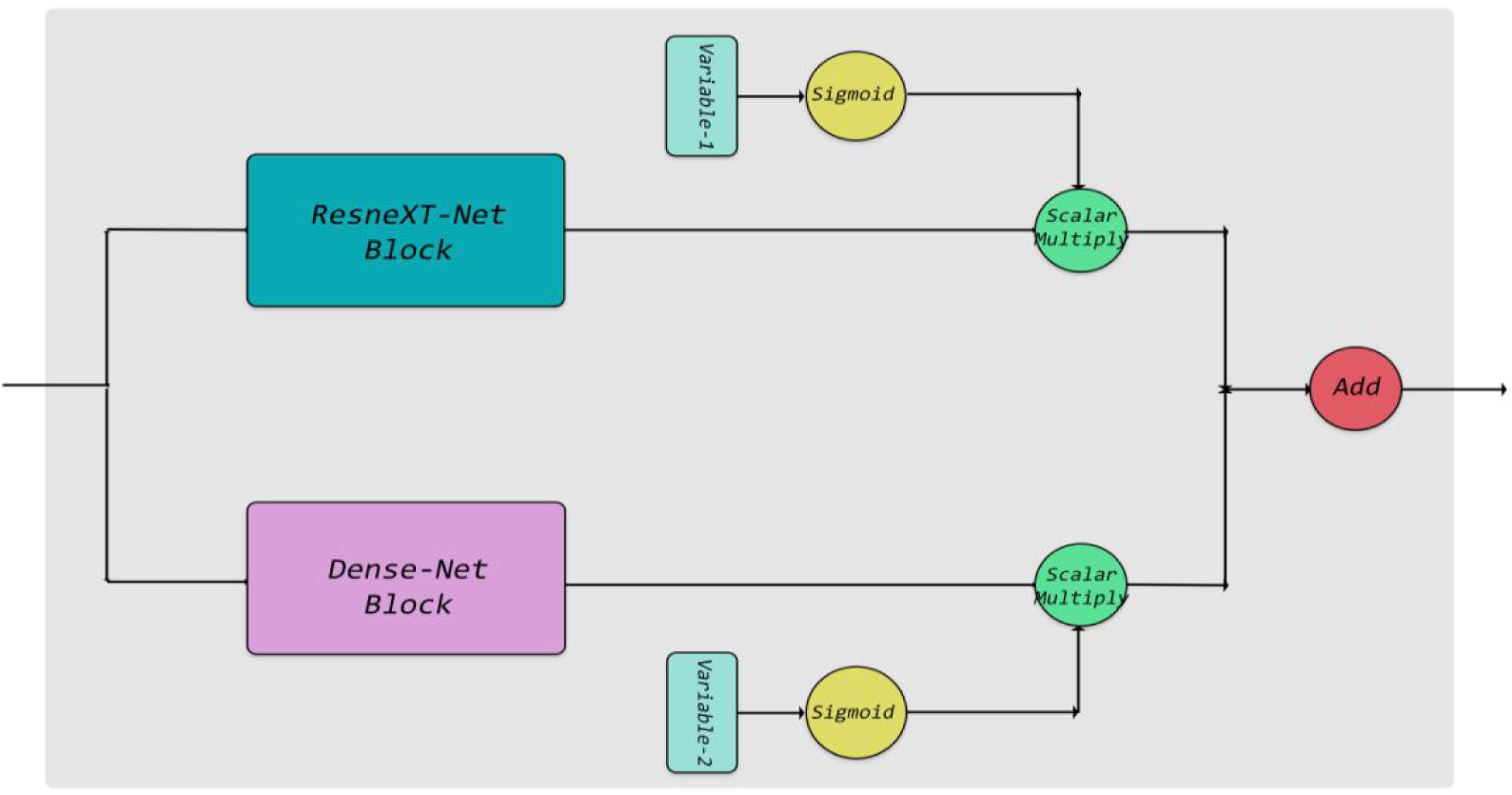
The structure of Combine Block.

### 8.6 Mimic Transformer Block

The mimic transformer block is the core of DCBTF net and the essential part of mimic transformer block is multi-head block. It composes with three parallel blocks which combine with a small-unit block and one combine block. The results of those three parallel blocks will be concated to one tensor. After that, the small-unit block which has 1×1 kernel will give an nonlinear transforming to this concated tensor. The add-normalization block is composed with an add operation and a layer normalization**[11]**. The Feed-Forward block is composed with 3 Small-Unit block and one Trans-Channels block. The kernels size of those 3 Small-Unit are 3×3, 2×2, 1×1 respectively. The kernel of Trans-Channels block is 1×1. The details of the structure of mimic transformer are at **Figure 24**.

**Figure 24:**
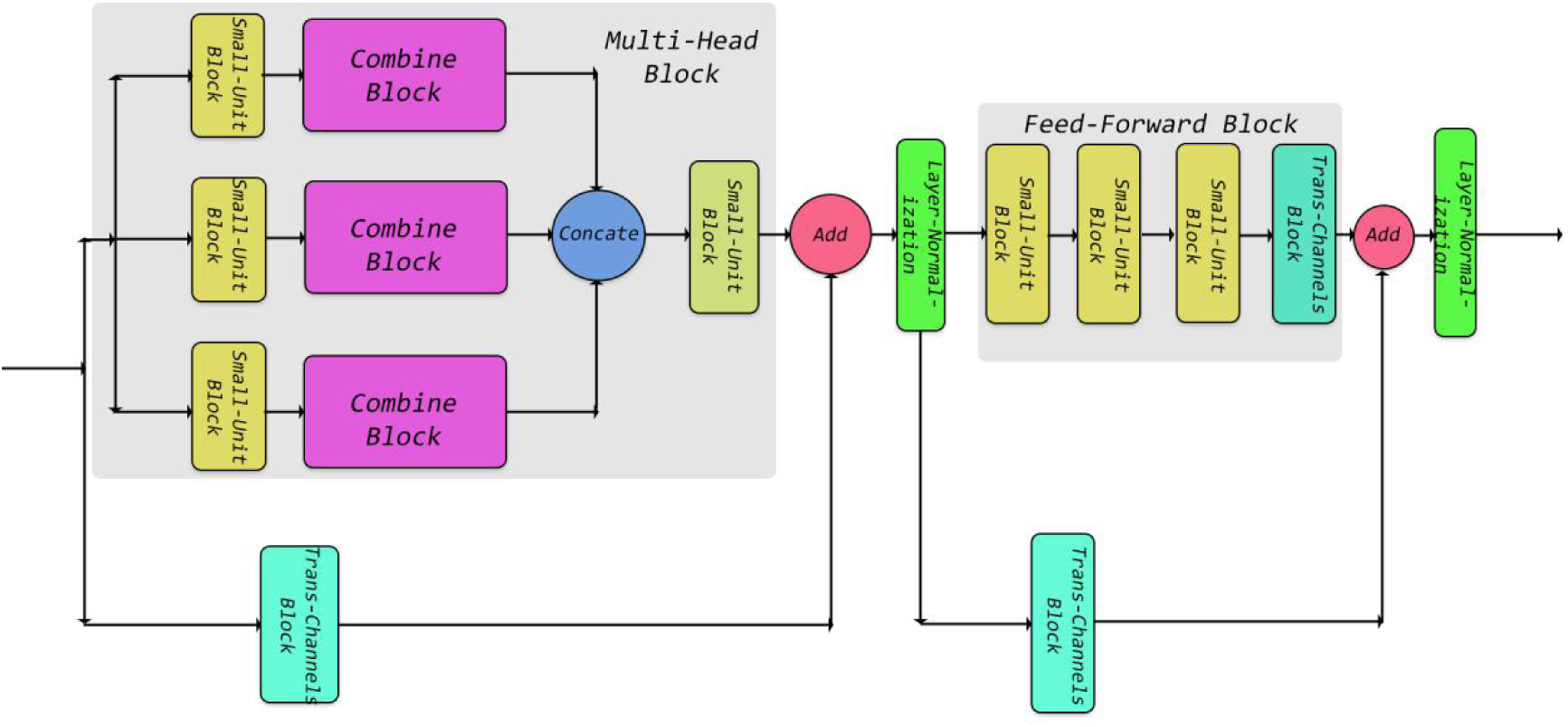
The structure of Mimic Transformer block.

### 8.7 Encoder Block

The encoder block composes with three parts, backbone network, feature pyramid network and multi-dimension predict with fusion information net. The details of structure of encoder are at **Figure 25**. The backbone network is used to extract high-level semantic feature. The feature pyramid network is used to merge high-level and low-level semantic information. The last part of encoder is used to predict result in different level and concats the results into one tensor. The mechanism of information gate is as same as the information gate in the combine block. It also can control how many information could pass to next layer. The residual block composes with a trans-channels block and one small-unit block with residual structure. In the residual block, the input tensor passes through small-unit block and trans-channels block simultaneously and add those results to one tensor. All of down or up sampling are sampled with 2 times.

**Figure 25:**
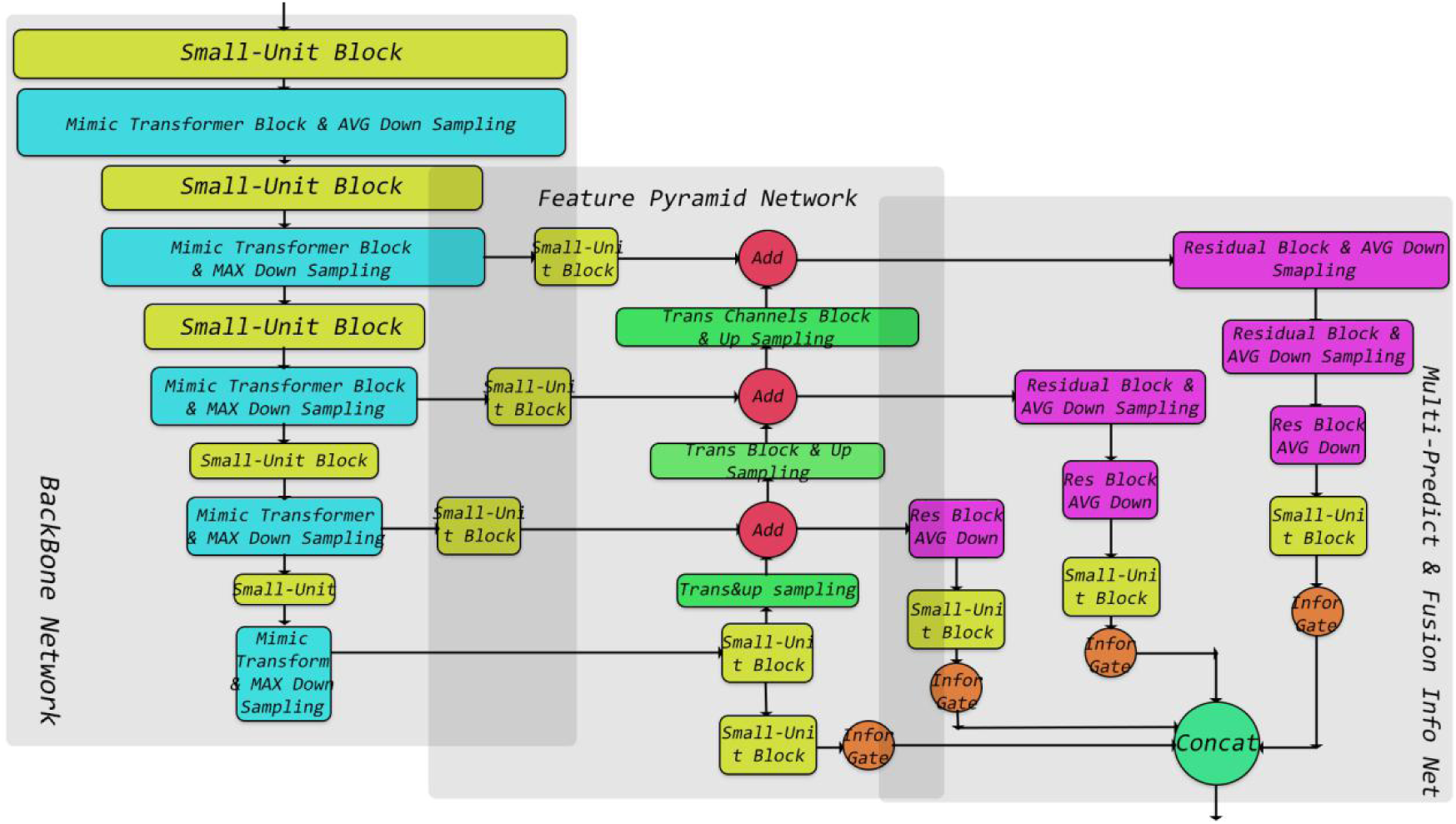
The detail of structure of encoder network.

### 8.8 Decoder Block

The decoder block is build by transpose blocks. There are double blocks and one transpose convolution operation in the transpose block and the residual structure is also appears in this block. The double blocks are small-unit and trans block respectively. The residual structure is similar residual block in encoder net. After adding double result, there is an transpose convolution for up sampling. For knowing the exact value of each pixel, i used transpose block five times for up sampling the high-level semantic feature to original size. The advantage of transpose convolution is that it contains variables and those variables can learn from data. There is an global average pooling at the bottom of the net to do down sampling. At the end, there is only one full connection without activation function to do classification.

## Footnotes

2 https://xenabrowser.net/datapages/?hub= https://tcga.xenahubs.net:443

3 https://xena.ucsc.edu/public-hubs/

4 The details of structure of DCBTF net are in the part of 8, Net Structure.

5 http://metascape.org/gp/index.html

